# Fibroblasts dynamically regulate lymphatic barrier function by modulating cell-cell junctions

**DOI:** 10.1101/2025.04.17.649442

**Authors:** Sarfaraz Ahmad Ejazi, Aisha Abdulkarimu, Theodore Hsiao, Rony Garcia-Vivas, Lina Berhaneyessus, Kathleen Trang, Alexander M. Xu, Katharina Maisel

## Abstract

Lymphatic dysfunction has been linked to several pathological conditions, including edema, inflammation, and cancer metastasis. The tight and adherens junction proteins between lymphatic endothelial cells (LECs) are important for preserving lymphatic vascular integrity. Despite the known role of fibroblasts in lymphangiogenesis, the direct impact of fibroblasts on LEC barrier function remains poorly understood. Here, normal human dermal fibroblast (NHDF) secretomes and non-contact co-culture models were used to examine how fibroblasts regulate human lymphatic endothelial cells (hLECs). Co-culture with dermal fibroblasts increased transendothelial electrical resistance (TEER) by > 110% and increased ZO-1 expression by >2-fold. VE-cadherin expression increased by at least 1.3-fold compared to controls. Furthermore, 4kDa dextran transport was markedly reduced, confirming an overall reduced hLEC permeability. Junction morphology analysis further showed a marked shift from discontinuous punctate and perpendicular junctions toward continuous ZO-1-rich borders, while actin anisotropy was reduced with redistribution away from aligned central stress fibers toward a more junction-supportive cytoskeletal architecture. Thrombin challenge revealed that fibroblasts also shape endothelial responses to inflammatory stress, indicating that stromal-endothelial crosstalk regulates both basal and stress-induced barrier behavior. Transcriptomic analysis identified two fibroblast-driven endothelial programs that converged on junctional remodeling but diverged in broader physiological features: activated NHDFs promoted lymphatic identity, junction stabilization, and a more quiescent endothelial state, whereas inactivated NHDFs favored extracellular matrix remodeling and partial EndoMT-like signaling while retaining some junction-supportive features. In contrast, lung fibroblast secretomes weakened barrier function, supporting the idea that stromal regulation of lymphatic permeability is tissue specific. Together, these findings identify fibroblasts as active regulators of lymphatic barrier physiology and reveal stromal heterogeneity as an important determinant of endothelial junction organization, transport, and tissue-specific lymphatic function.

**NEW AND NOTEWORTHY:** This exciting work demonstrates that fibroblasts are active regulators of lymphatic endothelial barrier physiology during homeostasis. Dermal fibroblasts strengthen barrier function and reduce paracellular transport by promoting continuous junction organization, cytoskeletal remodeling, and extracellular matrix reprograming, while fibroblast state and tissue origin produce distinct endothelial programs.

## INTRODUCTION

Lymphatic dysfunction is implicated in cardiovascular disease, fibrotic disease, chronic inflammation conditions, and cancer metastasis. Lymphatic vessels are distributed throughout most tissues and are essential for fluid homeostasis, immune cell trafficking, and dietary lipid absorption. Lymphatic vessels are lined by lymphatic endothelial cells (LECs), which adopt distinct structural and functional phenotypes along the lymphatic vascular tree. Initial lymphatics are specialized for uptake from peripheral tissues and are characterized by discontinuous button-like junctions, a sparse basement membrane, and the absence of perivascular cells. In contrast, collecting lymphatics are optimized for transport and propulsion of lymph and exhibit continuous zipper-like junctions, a more complete basement membrane, surrounding smooth muscle cells, and intraluminal valves that support directional flow.

Tight and adherens junctions are central regulators of lymphatic permeability and vessel integrity. Studies have highlighted the role of adherens junction proteins, including VE-cadherin, in supporting button-like junctions of initial lymphatics and disruption of VE-cadherin-dependent signaling increases lymphatic permeability and alters vessel function [1,2]. Tight junction proteins, including Claudin-5 and zonula occludin 1 (ZO-1), further regulate paracellular transport by organizing intercellular barrier structure and linking junctional complexes to the actin cytoskeleton. These junctions are dynamically regulated by inflammatory cytokines, growth factor signaling, matrix properties, and biomechanical forces such as shear stress, transmural flow, and interstitial pressure [3–7].

Advancements in lymphatic vascular research have highlighted the importance of the microenvironment in regulation of LEC phenotype [8]. In the interstitium, lymphatic vessels are embedded within extracellular matrix (ECM) and closely associated with stromal cells, immune cells, and perivascular support cells. Among these, fibroblasts are particularly well positioned to regulate lymphatic functions because they secrete ECM proteins, reshape tissue mechanics, and produce paracrine factors that influence endothelial signaling. Fibroblasts have already been shown to support lymphangiogenesis through VEGF-C and related pathways [9–12]. Additionally, fibroblast-associated signaling including FGF pathways can influence endothelial growth and sprouting [13].

In pathological settings, fibroblasts can also promote lymphatic remodeling and dysfunction; for example, cancer-associated fibroblasts enhance lymphangiogenesis, alter permeability, and contribute to metastatic spread [14,15]. Fibroblast-generated matrix has likewise been implicated in lymphatic regeneration during wound healing [16].

Most prior fibroblast-lymphatic co-culture studies have focused on lymphangiogenesis or immune and cancer cell trafficking in disease, leaving the homeostatic regulation of lymphatic barrier function largely unexplored. Here, we developed a non-contact in vitro co-culture model to define how fibroblasts regulate lymphatic cell-cell junctions, barrier integrity, and permeability. We first examined how secreted factors from normal human dermal fibroblasts (NHDFs) affect hLEC junction integrity and permeability and then studied fibroblast-LEC co-cultures on collagen beds, where NHDFs supported junction integrity and reduced paracellular transport. RNA sequencing further revealed fibroblast-driven transcriptional programs that regulate hLEC morphology and cytoskeletal organization in ways consistent with enhanced barrier function. Because stromal control of lymphatic permeability has direct implications for edema, immunotherapy delivery, and inflammation-driven lymphatic dysfunction, defining how fibroblasts tune lymphatic barrier properties during homeostasis may uncover mechanistic targets to restore lymphatic function in disease.

## RESULTS

### Exposure to fibroblast secretomes modulates lymphatic cell-cell junctions and reduces lymphatic permeability

To understand fibroblasts affect hLEC junctions, we first optimized the media conditions for fibroblast secretome collection. We found that the characteristic morphology and growth pattern of NHDF cells were similar in endothelial microvascular growth media (EGMV2), as this media also contains fibroblast growth factors. NHDFs were grown in either fibroblast growth media (FGM) with 1ng/mL of fibroblast growth factor and 5 µg/mL of insulin [NHDF(G)], or in basal FGM [NHDF(B)]. Then, NHDFs were exposed to EGMV2 low serum media for a day and NHDF secretome was collected. hLECs were grown as a monolayer on a transwell insert and then treated with NHDF(G) or NHDF(B) secretomes for 2 days and compared with hLEC cultured in EGMV2 media only without NHDF secretomes (hLEC alone) (**Figure 1A**).

**Figure 1.**
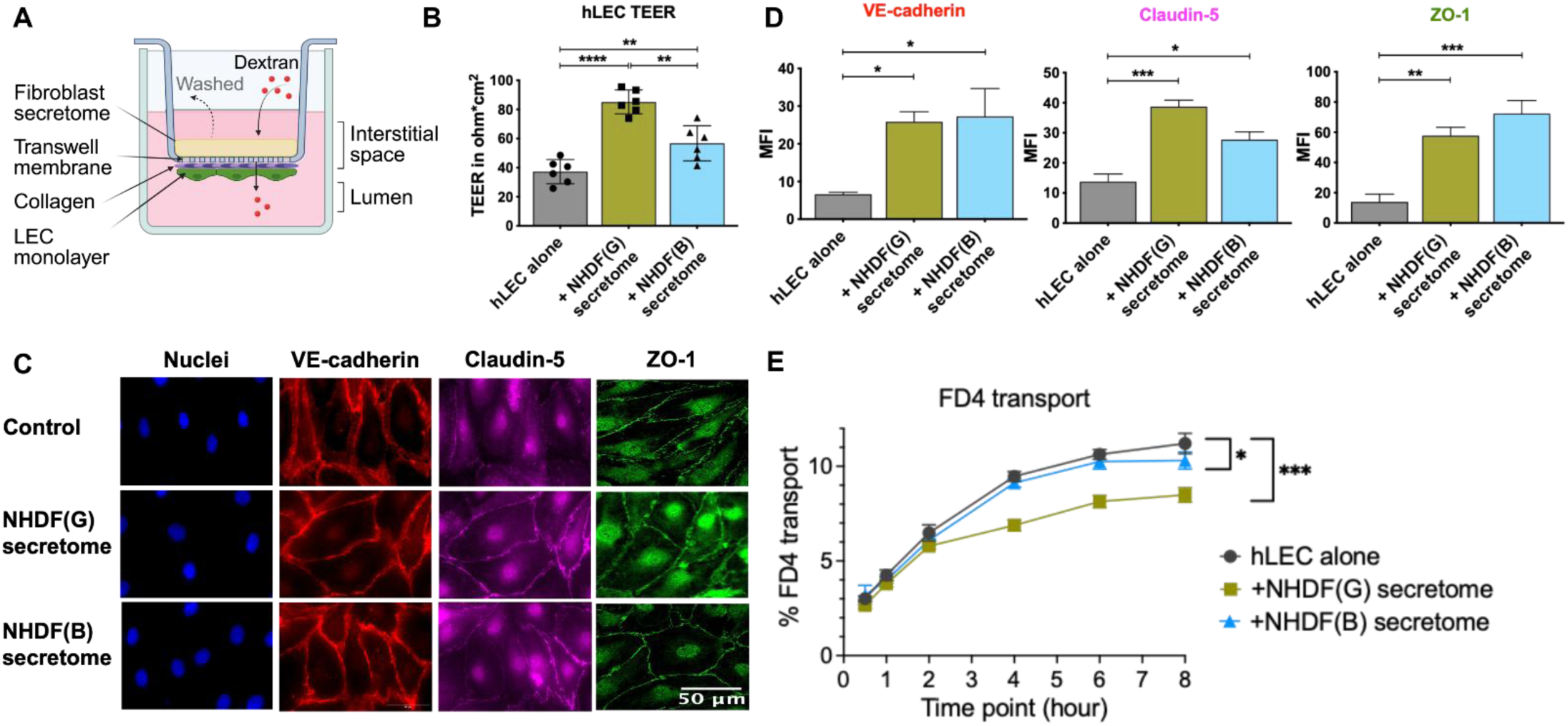
NHDF secretomes enhance hLEC barrier function. **(A)** Schematic representation of the transwell secretome assay. hLEC monolayers were treated with secretomes collected from activated (NHDF(G)) or inactivated (NHDF(B)) dermal fibroblasts for 48 hours. **(B)** Transendothelial electrical resistance (TEER) measurements of hLEC monolayers after treatment with NHDF(G) or NHDF(B) secretomes, compared with hLECs cultured alone in EGMV2 media. **(C)** Representative immunofluorescence images of VE-cadherin and Claudin-5, and confocal images of ZO-1 in hLECs following 48 h of secretome treatment. Nuclei were counterstained with Hoechst. Scale bar = 50 μm. **(D)** Quantification of VE-cadherin, Claudin-5, and ZO-1 fluorescence intensity using ImageJ (FIJI). **(E)** FITC-dextran (4 kDa) transport across hLEC monolayers following treatment with NHDF(G) or NHDF(B) secretomes. Data are presented as mean ± SEM from *n* = 6 independent experiments. Statistical significance in panels B and D was determined using one-way ANOVA followed by Dunnett’s multiple-comparisons test and Welch’s t-test. For the permeability assay (panel E), statistical significance was determined using two-way ANOVA followed by Tukey’s multiple-comparisons test (*n* = 3). ns, not significant (*P* ≥ 0.05); *P* < 0.05 (\**), P < 0.01 (*\*\**), P < 0.001 (*\*\*\*), and *P* < 0.0001 (****). Images were adjusted to improve visualization. Quantitative analyses were performed using the original unprocessed images.

We first assessed hLEC barrier integrity using transendothelial electrical resistance (TEER). Treatment with NHDF(G) and NHDF(B) secretomes resulted in significantly higher TEER values of 85 ± 8 Ω.cm^2^ and 57 ± 12 Ω.cm^2^, respectively, compared to 37 ± 8 Ω.cm^2^ for the hLEC alone control (**Figure 1B**). Next, we sought to analyze hLEC junction integrity and expression via immunofluorescence. We found that tight (ZO-1 and Claudin-5) and adherens (VE-cadherin) junction expression significantly increased with both the NHDF secretomes compared to hLEC alone in EGMV2 (**Figure 1C and 1D**). Specifically, treatment with NHDF(G) led to 4-, 3-, and 4-fold increases in the expression levels of VE-cadherin, Claudin-5, and ZO-1, respectively. NHDF(B) treatment resulted in 4-, 2-, and 5-fold upregulation of the junction proteins, respectively (**Figure 1D**). We also found that lymphatic endothelial markers, LYVE-1 and PROX1, were not altered in fluorescence intensity (**Supplementary Figure 1A and 1B**). To evaluate changes in hLEC monolayer permeability following treatment with fibroblast secretomes, we assessed transport of a fluorescently labeled, 4kDa dextran (FD4). We observed a 24% reduction in permeability, decreasing from 11 ± 1% in the hLEC alone group to 8 ± 1% with NHDF(G) secretome-treated group after 8 hours (**Figure 1E**). These findings indicate that dermal fibroblasts’ secretome promotes low-permeability phenotype in hLECs, which may explain how dermal lymphatics remain resistant to leakage even under elevated interstitial fluid pressure.

### Co-culture with fibroblasts regulates LEC junction and reduces lymphatic permeability

To address the direct effects of fibroblasts on the lymphatic junction expression and permeability, we generated a non-contact co-culture model (**Figure 2A**), where hLECs are cultured on the bottom of the transwell and after monolayers were established, NHDFs were seeded on the transwell top. Cells were co-cultured for 2 days, after which fibroblasts were scraped from the membrane to prevent interference with TEER and permeability measurements (**Figure 2B**). hLEC monolayers exhibited increased TEER following co-culture with NHDF(G) and NHDF(B), rising from 46 ± 13 Ω.cm^2^ in hLECs cultured alone to 112 ± 5 Ω.cm^2^ and 95 ± 15 Ω.cm^2^, respectively (**Figure 2C**). We also found a distinct increase in ZO-1 and VE-cadherin expression in both hLECs co-cultured groups (+NHDFs) (**Figure 2D**). VE-cadherin expression increased by 1.6- and 1.3-fold, whereas ZO-1 expression increased by 2.1- and 2.0-fold in hLECs co-cultured with NHDF(G) and NHDF(B), respectively, compared with hLECs cultured alone (**Figure 2E**). FD4 permeability across the hLEC monolayer decreased by 24% following co-culture with NHDF(G), and by 9.5% with NHDF(B), indicating reduced permeability (**Figure 2F)**.

**Figure 2.**
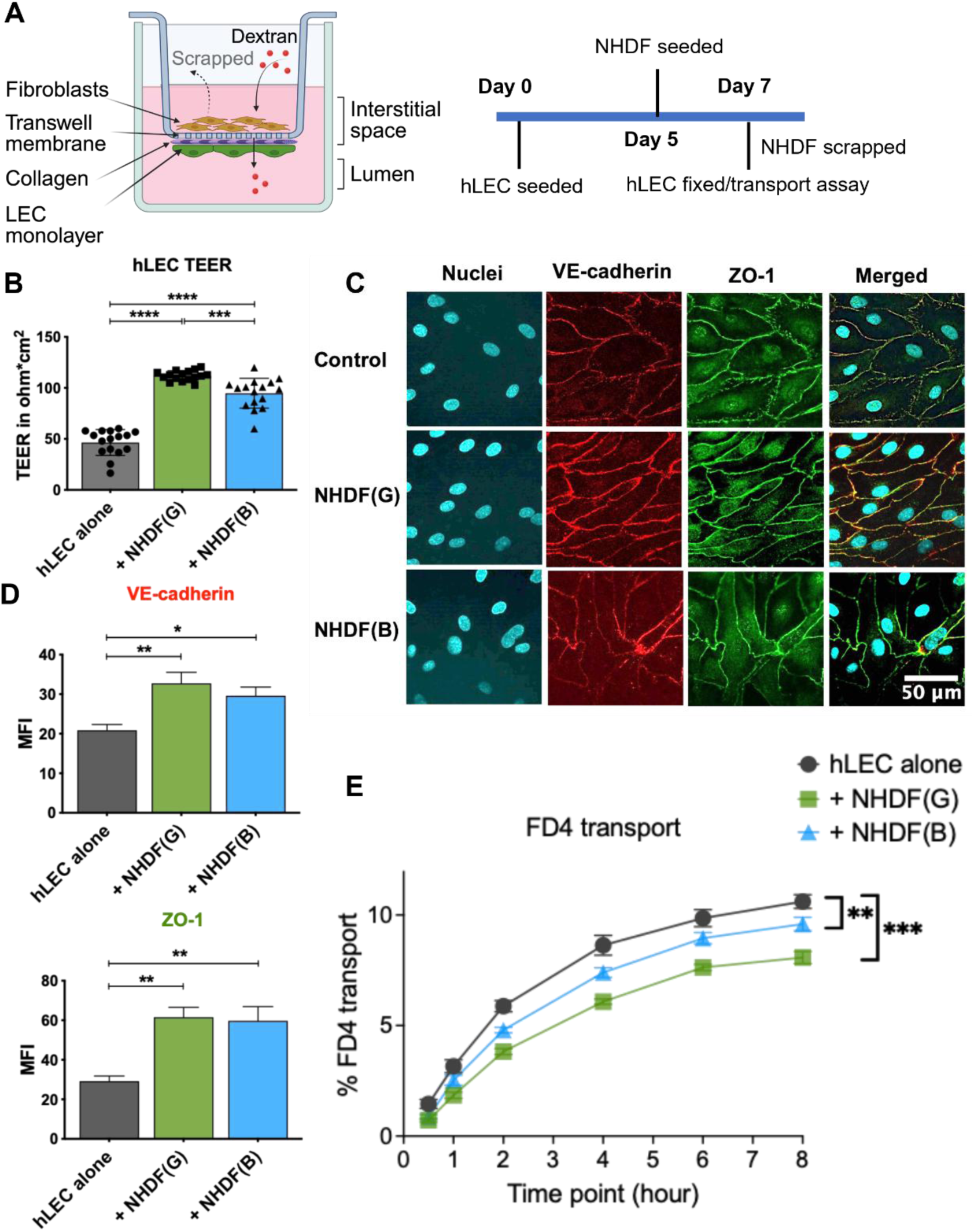
NHDF co-culture enhances hLEC barrier function. **(A)** Schematic representation of the transwell co-culture assay and **(B)** experimental timeline. hLECs were cultured with activated (NHDF(G)) or inactivated (NHDF(B)) dermal fibroblasts for 48 hours after which the fibroblasts were removed prior to downstream analyses. **(C)** Transendothelial electrical resistance (TEER) was measured across hLEC monolayers following 48 hours of co-culture with NHDF(G) or NHDF(B), compared with hLECs cultured alone. TEER reflects the barrier function of the hLEC monolayer. **(D)** Representative confocal images of VE-cadherin and ZO-1 in hLECs following 48 h of co-culture with NHDF(G) or NHDF(B). Nuclei were counterstained with Hoechst. Scale bar = 50 μm. **(E)** Quantification of VE-cadherin and ZO-1 fluorescence intensity using FIJI/ImageJ. **(F)** FITC-dextran (4 kDa) transport across hLEC monolayers following co-culture with NHDF(G) or NHDF(B). Data are presented as mean ± SEM from *n* = 6 independent experiments. Statistical significance in panels B and D was determined using one-way ANOVA followed by Dunnett’s multiple-comparisons test and Welch’s t-test. For the permeability assay (panel E), statistical significance was determined using two-way ANOVA followed by Tukey’s multiple-comparisons test (*n* = 9). ns, not significant (*P* ≥ 0.05); *P* < 0.05 (\**), P < 0.01 (*\*\**), P < 0.001 (*\*\*\*), and *P* < 0.0001 (****). Images were adjusted to improve visualization. Quantitative analyses were performed using the original unprocessed images.

We performed detailed analysis of LEC junction morphology using the Junction Analyzer Program (JAnaP) [17], a Python-based program that quantitatively analyzes cell-cell junction presentation along with cell morphology. We found no difference in the cellular solidity but did see an 18% decrease in the cell circularity of hLECs co-cultured with NHDF(G) (**Figure 3A**). Both NHDF(G) and NHDF(B) promote lymphatic monolayers, increasing hLEC coverage by 7%. We also found a significant increase in ZO-1 continuous junctions from 84 ± 5% in hLEC alone to 96 ± 2% in both co-cultures. Similarly, both co-cultures significantly reduced ZO-1 discontinuous junctions, with reductions in punctate junctions by 89% and 97% and perpendicular junctions by 64% and 62% with NHDF(G) and NHDF(B), respectively (**Figure 3A**). **Figure 3B** illustrates the criteria used by the Junction Analyzer Program to categorize junctional morphology.

**Figure 3.**
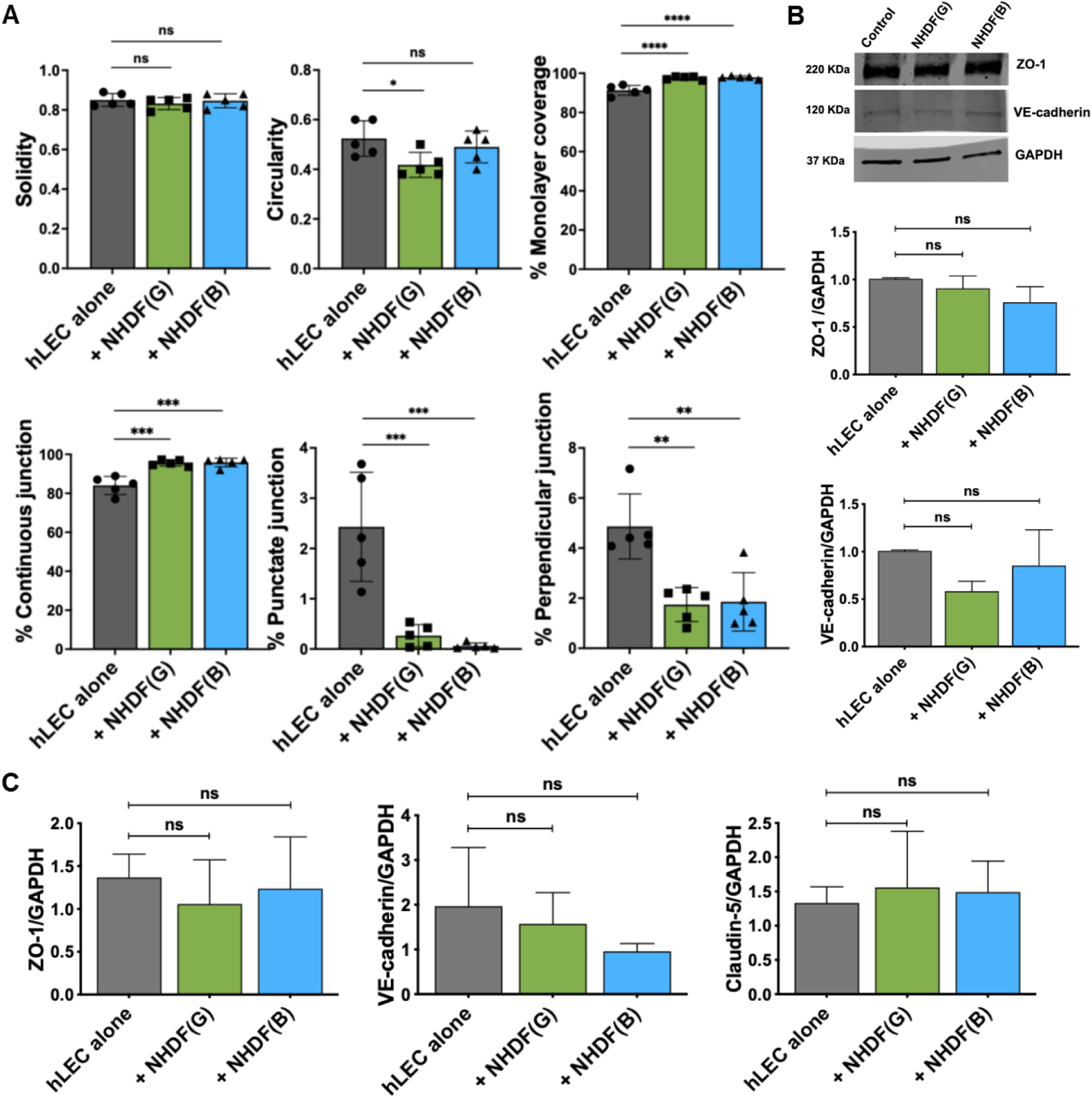
NHDF co-culture promotes hLEC junctional organization. **(A)** Quantitative analysis of hLEC junction morphology and organization for ZO-1 immunofluorescence using JAnaP following 48 hours of co-culture with NHDF(G) or NHDF(B). Parameters analyzed included cell solidity, circularity, monolayer coverage, and the proportions of continuous, punctate, and perpendicular junctions. **(B)** Schematic representation of the junctional analyzer program algorithm used to categorize continuous, punctate, and perpendicular junctions. **(C)** Immunoblot and densitometric quantification of ZO-1 (220 kDa) and VE-cadherin (120 kDa) protein expression in hLECs following co-culture with NHDF(G) or NHDF(B). GAPDH was used as a loading control. **(D)** RT-qPCR analysis of ZO-1, VE-cadherin, and Claudin-5 mRNA expression levels in hLECs under the indicated conditions. Data are presented as mean ± SEM from independent experiments. Statistical significance was assessed using one-way ANOVA followed by Dunnett’s multiple-comparisons test. ns, not significant (*P* ≥ 0.05); *P* < 0.05 (\**), P < 0.01 (*\*\**), P < 0.001 (*\*\*\*), and *P* < 0.0001 (****).

Immunoblot analysis showed no notable increase in overall ZO-1 (220 kDa) and VE-cadherin (120 kDa) protein levels in co-cultured hLECs compared to the control (**Figure 3C**). Similarly, mRNA expression levels of ZO-1, VE-cadherin and Claudin-5 (**Figure 3D**), along with lymphatic markers, PROX-1 and LYVE-1 (**Supplementary figure 1C**) did not differ significantly from the control. These findings confirm our previous observation that fibroblast presence alone confers a barrier enhancement, though the underlying mechanisms remain to be determined.

### Fibroblast modulates actin cytoskeletal organization in LECs

Since fibroblast co-culture induced changes in endothelial cell morphology and junctional protein expression, we next examined intracellular F-actin organization. Actin filament alignment was quantitatively assessed from immunofluorescence images using the FibrilTool plugin in ImageJ (FIJI), which measures fiber anisotropy. hLECs cultured alone exhibited a moderately aligned actin cytoskeleton, with an average anisotropy of ∼0.22. In contrast, hLECs co-cultured with NHDF(G) displayed a marked reduction in actin alignment, with anisotropy values decreasing approximately five-fold (∼0.04) (**Figure 4A**). Morphologically, F-actin organization in this condition appeared less structured, with reduced central stress fiber alignment and a redistribution of actin toward junction-associated regions (**Figure 4B**). hLECs co-cultured with NHDF(B) exhibited an intermediate phenotype, with partial preservation of actin organization. Anisotropy values (∼0.07) were reduced 3-fold compared to control but remained higher than those observed in NHDF(G). Notably, F-actin in NHDF(B)-conditioned hLECs showed a predominantly cortical arrangement, with actin fibers enriched along the cell periphery. Consistent with the co-culture findings, treatment with NHDF(G) and NHDF(B) secretomes produced comparable reductions in actin anisotropy **(Figures 4C)** and stress fiber alignment **(Figure 4D)**. These findings indicate that fibroblast conditioning remodels F-actin organization, reducing stress fiber alignment and promoting a more cortical actin network. This cytoskeletal reorganization was associated with improved junctional integrity and reduced permeability, suggesting a close relationship between actin organization and hLEC barrier function.

**Figure 4.**
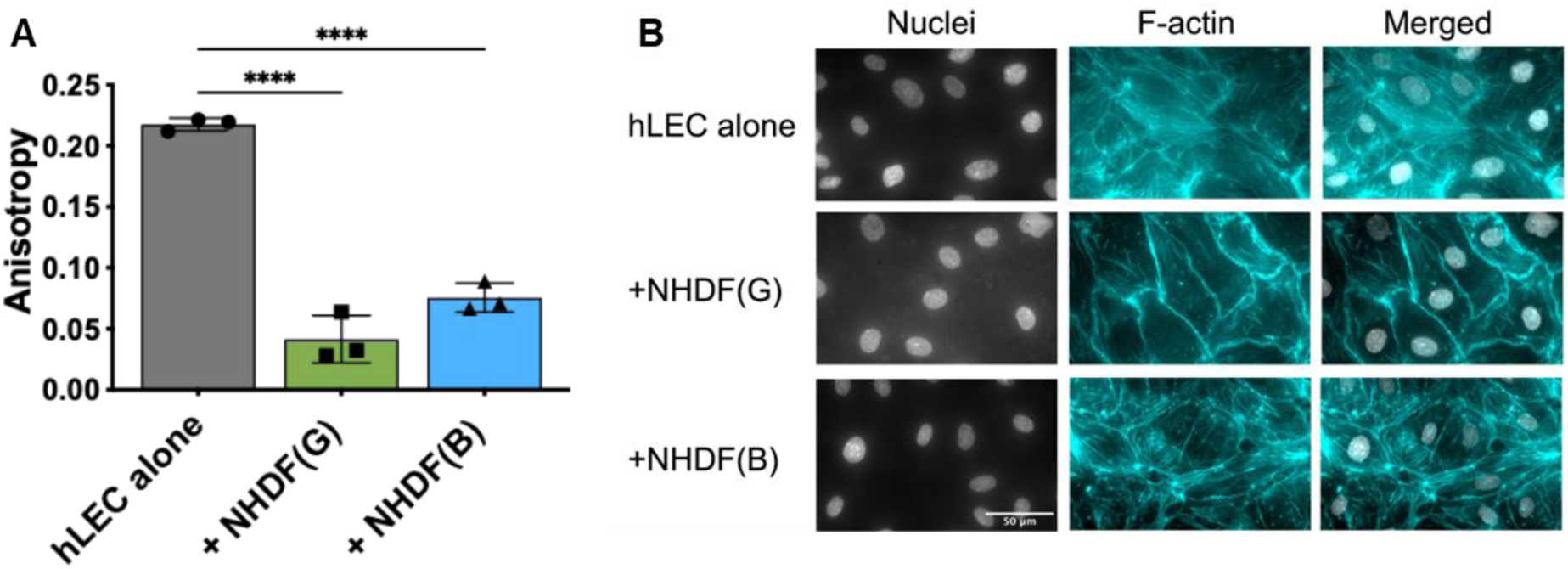
Fibroblast remodels actin organization in hLECs. **(A)** Quantification of actin fiber alignment in hLECs co-cultured with NHDFs or **(C)** conditioned with NHDFs secretomes. Actin anisotropy was measured using the FibrilTool plugin in ImageJ (FIJI). Anisotropy values range from 0 to 1, with higher values indicating greater actin filament alignment. At least 15 or 9 cells per condition from three independent biological replicates were analyzed. Statistical significance was determined using one-way ANOVA followed by Dunnett’s multiple-comparisons test. ns, not significant (*P* ≥ 0.05); *P* < 0.05 (\**), P < 0.01 (*\*\**), P < 0.001 (*\*\*\*), and *P* < 0.0001 (****). **(B)** Representative immunofluorescence images of F-actin organization in hLECs co-cultured with NHDFs or **(D)** conditioned with NHDFs secretomes. Nuclei are shown in grayscale and F-actin in cyan. Merged images are shown in the right column. Scale bar = 50 μm. Images were adjusted to improve visualization. Anisotropy was performed using the original unprocessed images.

### Fibroblasts promote thrombin-induced LEC barrier dysfunction

We next investigated whether fibroblast conditioning could reverse disruption of LEC barrier function by thrombin. Thrombin, a potent pro-inflammatory mediator, disrupts endothelial barrier integrity leading to vascular dysfunction and increase endothelial permeability in the blood vasculature [18]. While thrombin clotting can occur in lymphatic vessels, the effects of thrombin signaling on lymphatic function, particularly its impact on permeability, remain to be fully understood. We examined the impact of thrombin on hLECs in the presence of fibroblast, NHDF(G). hLECs were either exposed to thrombin for 4 hours or left untreated, followed by a 48 h co-culture with NHDF. Thrombin treatment caused a 19% reduction in TEER in hLECs alone, and a 12% reduction in LECs co-cultured with NHDF(G) (**Figure 5A**). Thrombin treatment also reduced hLEC numbers in both the hLEC alone and NHDF(G) co-culture group, suggesting a potential impact on cell viability or proliferation (**Figure 5B**). Additionally, we assessed the permeability of hLECs to FD4 after thrombin treatment. Consistent with the TEER findings, thrombin increased dextran transport by 30% in hLECs in the presence of NHDF(G) (**Figure 5C**). Immunofluorescence of junction proteins in thrombin-treated hLECs showed an irregular junction protein expression. When hLECs were co-cultured with NHDF(G) after thrombin treatment, they exhibited noticeably higher junction discontinuity (**Figure 5D**). These results indicate that fibroblast-conditioned hLECs remain highly susceptible to acute thrombin challenge and exhibit enhanced thrombin-induced disruption, highlighting stromal-endothelial crosstalk as a critical regulator of thrombo-inflammatory dysfunction. We next sought to define the specific cellular processes transcriptionally regulated by fibroblast conditioning and the mechanisms driving that regulation.

**Figure 5.**
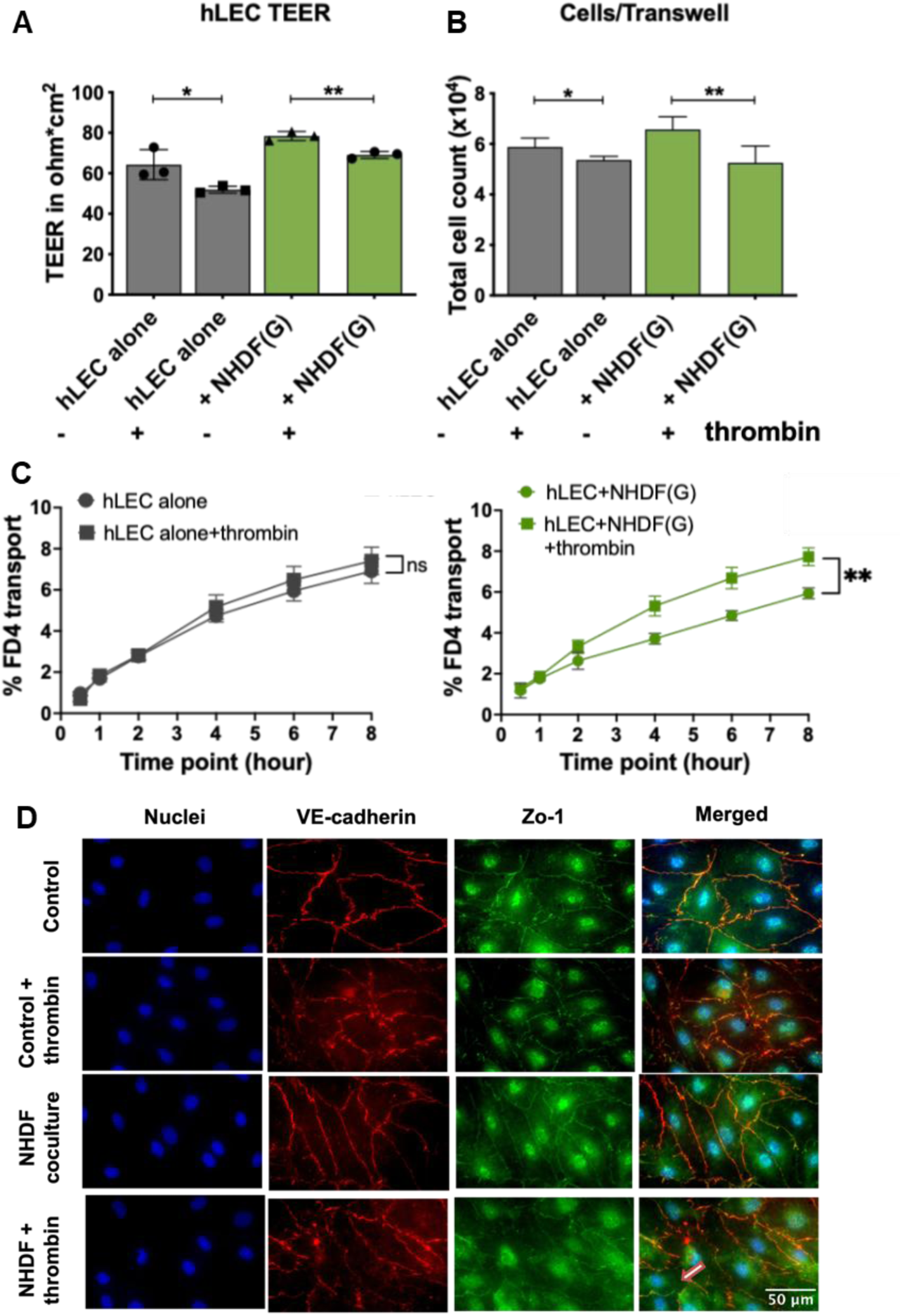
Thrombin attenuates fibroblast-mediated enhancement of hLEC barrier function. **(A)** Transendothelial electrical resistance (TEER) measurements of hLEC monolayers cultured alone or following 48 hours of co-culture with NHDF(G), with or without 4 hours of thrombin treatment (*n* = 3). **(B)** Total hLEC counts per transwell under the indicated conditions (*n* = 6). **(C)** FITC-dextran (4 kDa) transport across hLEC monolayers following 48 hours of NHDF(G) co-culture and/or 4 hours of thrombin treatment. **(D)** Representative immunofluorescence images of VE-cadherin (red) and ZO-1 (green) in hLECs cultured alone or following NHDF(G) co-culture, with or without thrombin treatment. Nuclei were counterstained with Hoechst (blue). Scale bar = 50 μm. Data are presented as mean ± SEM from 3–6 independent experiments. Statistical significance in panels A and B was determined using unpaired two-tailed Student’s t-tests comparing thrombin-treated and untreated wells within each culture condition. For the transport assay (panel C), statistical significance was determined using two-way ANOVA followed by Tukey’s multiple-comparisons test. ns, not significant (*P* ≥ 0.05); *P* < 0.05 (\**), P < 0.01 (*\*\**), P < 0.001 (*\*\*\*), and *P* < 0.0001 (****). Images were adjusted to improve visualization. Quantitative analyses were performed using the original unprocessed images.

### RNA-seq identifies fibroblast-derived reprogramming of lymphatic endothelial junctions across distinct pathways

To examine fibroblast-mediated regulation of lymphatic endothelial phenotype, bulk RNA-seq was performed. Significant differentially expressed genes (DEGs) were identified between hLECs cultured alone and hLECs co-cultured with NHDF(G) and NHDF(B) for 48 h, revealing eight major molecular mechanisms contributing to fibroblast-mediated regulation of lymphatic endothelial phenotype **(Table 1)**. NHDF(G)-conditioned hLECs displayed a transcriptional signature associated with junction remodeling, lymphatic maturation, and immune activation, whereas NHDF(B)-conditioned hLECs exhibited a transcriptional profile enriched in ECM remodeling, cytoskeletal reprogramming, and endothelial-to-mesenchymal transition (EndoMT)-related processes **(Figure 6A) (Supplementary Figure 2).**

**Figure 6.**
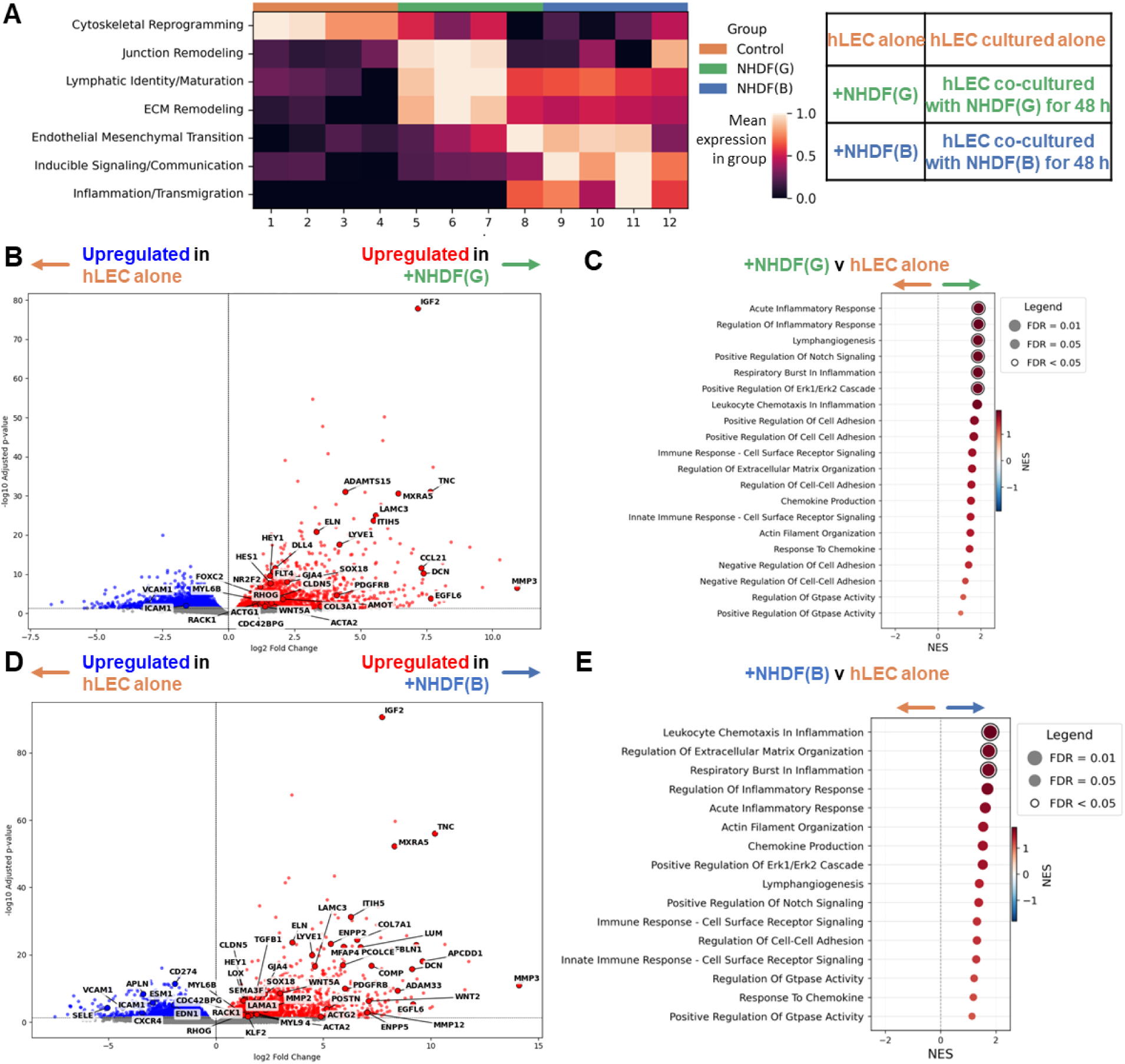
Fibroblast-induced transcriptomic remodeling of hLECs. **(A)** Heatmap showing the gene set scoring (expression) of representative genes associated with major biological pathways in hLECs cultured alone or following 48 hours of co-culture with NHDF(G) or NHDF(B). The inset illustrates the experimental groups. **(B)** Volcano plots of differentially expressed genes (DEGs) in hLECs following NHDF(G) or **(D)** NHDF(B) co-culture, respectively, compared with hLECs cultured alone. Red dots represent significantly upregulated genes (adjusted p < 0.05 and log_2_ fold change > 1), whereas blue dots represent significantly downregulated genes (adjusted p < 0.05 and log_2_ fold change < −1). **(C)** Gene set enrichment analysis (GSEA) dot plots for hLECs following NHDF(G) or **(E)** NHDF(B) co-culture, respectively, compared with hLECs cultured alone. Pathways pertaining to regulation of biological processes contain both positive and negative regulation responses. Positively enriched pathways are shown in red, whereas negatively enriched pathways are shown in blue. Dot color represents the normalized enrichment score (NES), and dot size corresponds to – log_10_ (FDR q-value). Fibroblasts were removed before RNA isolation, and all transcriptomic analyses were performed exclusively on hLECs.

**Table 1.** Representative RNA-seq-based differentially expressed genes (DEGs). DEGs associated with key biological pathways in hLECs cultured alone or following 48 hours of co-culture with NHDF(G) or NHDF(B). Underlined genes are unique to the indicated comparison. Green indicates upregulated genes, and red indicates downregulated genes.

| Transcriptomic response | +NHDF(G) v hLEC alone | +NHDF(B) v hLEC alone | +NHDF(B) v +NHDF(G) |
| --- | --- | --- | --- |
| **Junctional remodeling** | CLND5, GJA4, <u>AMOT</u> | CLND5, GJA4, <u>JAM2</u>, <u>CTNNB1</u> |  |
| **Cytoskeletal reprogramming** | RHOG, CDC42BPG, RACK1, MYL6B, KLF2, TGFB1, <u>ACTG1</u>, RAC3 | RHOG, CDC42BPG, RACK1, MYL6B, <u>MYL9</u>, <u>ACTG2</u>, <u>TAGLN</u>, KLF2, TGFB1 |  |
| **Lymphatic identity/maturation** | LYVE-1, <u>CCL21</u>, <u>FLT4</u>, <u>FOXC2</u>, <u>NR2F2</u>, SOX18, DLL4, <u>HES1</u>, HEY1, STAB2, WNT5A, APCDD1, VEGFB | LYVE-1, SOX18, SEMA3F, HEY1, WNT5A, VEGFB, VEGFC, <u>CXCR4</u>, <u>APLN</u> | <u>CCL21</u>, <u>HES1</u>, <u>CXCR4</u>, <u>STAB2</u>, <u>CLEC4M</u>, <u>APLN</u> |
| **Inflammation/transmigration** | <u>VCAM1</u>, <u>ICAM1</u>, <u>SELE</u>, <u>EDN1</u>, <u>ESM1</u>, <u>PD-L1</u> | <u>VCAM1</u>, <u>ICAM1</u>, <u>SELE</u>, <u>EDN1</u>, <u>ESM1</u>, <u>PD-L1</u> |  |
| **ECM remodeling** | MMP3, IGF2, TNC, DCN, LOXL1, LAMC3, ITIH5, ADAMTS15, ELN, MXRA5, EGFL6, LUM, COMP, <u>ITGB1</u> | MMP3, <u>MMP2</u>, <u>MMP12</u>, IGF2, TNC, DCN, LAMC3, <u>LAMA1</u>, ITIH5, ADAM33, ELN, MXRA5, EGFL6, LUM, COMP, <u>POSTN</u>, <u>FBLN1</u>, <u>COL7A1</u>, <u>PCOLCE</u>, <u>LOX</u>, <u>MFAP4</u>, <u>ENPP2</u>, <u>ENPP5</u>, <u>ITGA9</u>, <u>ITGB1</u> | COMP, MMP3, MMP12, TNC, DCN, LAMA1, GPNMB, HGF, THBS2, <u>ITGA9</u> |
| **Endothelial-to-mesenchymal transition** | <u>COL3A1</u>, ACTA2, <u>ACTG1</u>, PDGFRB, WNT4 | ACTA2, <u>ACTG2</u>, PDGFRB, WNT4, <u>WNT2</u>, APCDD1, <u>THY1</u> | PDGFRA, <u>WNT4</u> |
| **Immune activation** | LILRB1, LCP2, ZAP70, HAVCR2, CRAD11, CD53, | <u>LAMP5</u>, <u>LAMP3</u> | <u>LILRB1</u>, <u>LCP2</u>, <u>ZAP70</u>, <u>HAVCR2</u>, <u>CARD11</u>, <u>CD53</u>, <u>LAMP5</u> |
| **WNT signaling** | FRZB, GREM2 | RSPO3, WNT5A, WNT2, FRZB, SFRP4, GREM1, GREM2 | RSPO3, WNT5A, FRZB, SFRP4, GREM1, GREM2 |

### NHDF(G)-conditioning promotes lymphatic junction remodeling and lymphatic maturation

Direct comparison between NHDF(G) co-cultured hLECs and hLECs co-cultured alone identified an upregulation of genes associated with lymphatic identity (e.g. LYVE-1, FOXC2), maintenance of junction integrity (e.g. CLDN5), actin dynamics, cytoskeletal remodeling (e.g. CDC42BPG, RHOG), and ECM remodeling (e.g. IGF2 and MMP3) (**Figure 6B).** NHDF(G)-co-cultured hLECs regulate genes associated with actin dynamics and cytoskeletal remodeling, which triggers a cascade that ultimately maintains lymphatic junctional integrity. This is evidenced by the upregulation of the tight junction protein CLDN5, the gap junction protein GJA4, and AMOT, a key regulator linking the cytoskeleton to cell-cell junctions. This is paired with reduced inflammatory activation indicative of a tighter LEC barrier formation and downregulation of genes involved in leukocyte adhesion and transmigration (i.e. VCAM1, ICAM1) are downregulated (**Figure 6B**). Consistently, Gene Set Enrichment Analysis (GSEA) showed significant enrichment of inflammatory response regulation, Notch and Erk1/Erk2 signaling pathways and lymphangiogenesis in NHDF(G) conditioned hLECs compared to the hLECs alone. Positive enrichment trends were found for junction assembly, cell-cell adhesion and GTPase activity (**Figure 6C**). These findings indicate that hLECs cultured under NHDF(G) conditions support a dynamic, coordinated enhancement of lymphatic identity and junctional stability, thereby regulating junction presentation at the hLEC perimeter.

### NHDF(B) enhances ECM remodeling and mesenchymal transition of lymphatic endothelial cells

Comparison of NHDF(B)-co-cultured hLECs with hLECs cultured alone revealed significant enrichment of ECM remodeling genes (**Figure 6D**). This includes matrix-degrading enzymes (e.g. MMP3, MMP2, MMP12), regulators of matrix stiffness and elasticity (e.g. LOX, elastin), proteins associated with fibrotic remodeling and matrix plasticity (e.g. POSTN, TNC), and ECM-associated signaling mediators (e.g. HGF, ENPP2, IGF2, EGFL6) (**Figure 6D**). NHDF(B) co-culture markedly upregulated EndoMT-associated genes (e.g. ACTA2 (α-SMA), PDGFRB) upstream signaling drivers (e.g. WNT4 and WNT2) and transition modulators (e.g. THY1, APCDD1). GSEA further highlighted significant enrichment in the regulation of the inflammatory response and ECM regulation pathways. We additionally identified positive enrichment trends for lymphangiogenesis and cell-cell adhesion pathway which suggest broader regulatory changes in these processes (**Figure 6E**). We then asked which DEGs were shared or differed between hLEC co-cultured conditioning between NHDF(G) and NHDF(B).

### Distinct transcriptional responses between hLECs co-cultured with NHDF(G) and NHDF(B) in ECM remodeling and immune activation

Direct comparison of the NHDF(G)- and NHDF(B)-conditioned hLECs revealed distinct regulatory programs that shape the lymphatic endothelial phenotype (**Figure 7 and Table 1**). NHDF(G) induced a transcriptional shift toward lymphatic identity and immune activation, including upregulation of CCL21, CLEC4M, CXCR4, LAMP5, and LILRB1 (**Figure 7A).** NHDF(G) also induced a transcriptional shift away from ECM remodeling, supporting ECM stabilization, as evidenced by downregulation of MMP3, MMP12, and TNC **(Figure 7A).** This is further supported by GSEA analysis, which indicates a depletion in collagen fibril organization (**Figure 7B**). The presence of NHDF(G) enforces hLEC junction stability through enrichment of cell-cell junction assembly, cell-cell adhesion, and lymphangiogenesis whereas NHDF(B) associates with ECM reorganization and mesenchymal transition. While 2,839 genes were commonly regulated, 1,562 and 1,357 genes were uniquely altered in the NHDF(G)- and NHDF(B)-conditioned hLECs (**Figure 7C)**.

**Figure 7.**
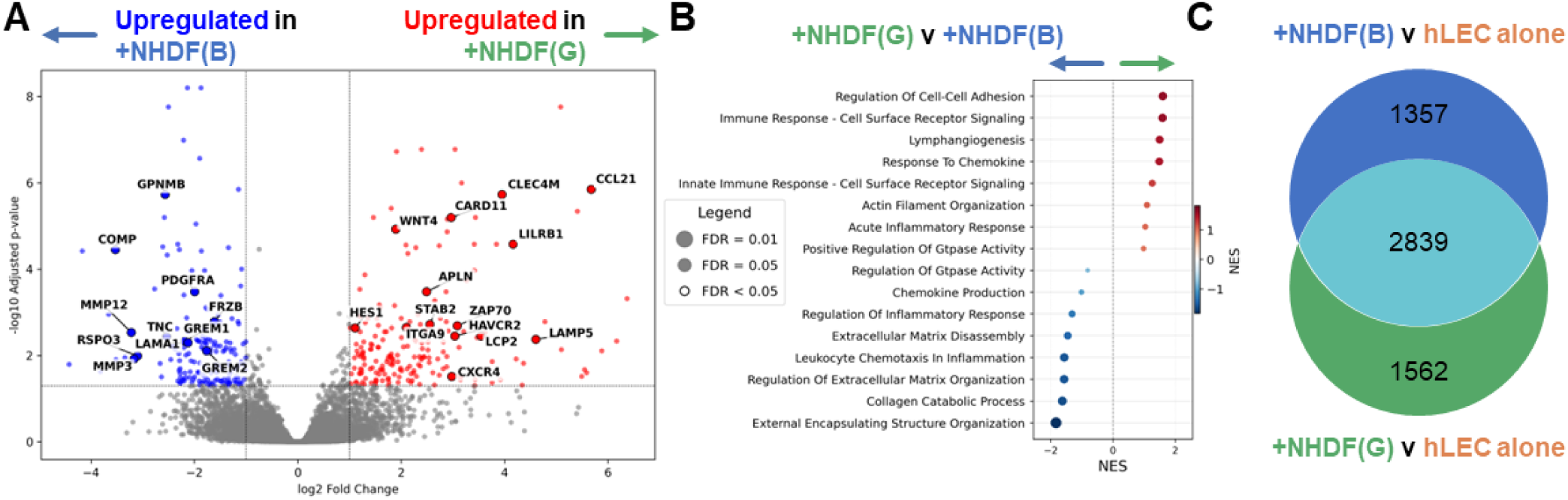
hLECs co-cultured with NHDF(G) or NHDF(B) revealed distinct regulatory programs. **(A)** Volcano plot showing differentially expressed genes (DEGs) between NHDF(G)- and NHDF(B)-co-cultured hLECs. Genes with an adjusted *p*-value < 0.05 and an absolute log_2_ fold change > 1 are highlighted, with selected representative genes annotated. Red and blue dots indicate significantly upregulated and downregulated genes, respectively, in NHDF(G) relative to NHDF(B), while gray dots represent non-significant genes. **(B)** Gene Set Enrichment Analysis (GSEA) dot plot comparing NHDF(G) and NHDF(B) co-culture conditions. Pathways pertaining to regulation of biological processes contain both positive and negative regulation responses. Positively enriched pathways in NHDF(G)-co-cultured hLECs are shown in red, whereas pathways enriched in NHDF(B)-co-cultured hLECs are shown in blue. Dot color represents the normalized enrichment score (NES), and dot size corresponds to – log_10_ (FDR q-value). **(C)** Venn diagram illustrating the overlap of significant DEGs (|log_2_ fold change| > 1, adjusted *p*-value < 0.05) identified in NHDF(B) versus control and NHDF(G) versus control comparisons.

Taken together, these findings indicate that the biochemical composition of the perivascular and interstitial tissues is crucial in regulating key cellular phenomena and processes that guide and regulate lymphatic function. Our study suggests that fibroblasts modulate hLEC function regardless of their activation status. We then asked whether the effect we are seeing is tissue-specific or whether fibroblast secretomes derived from different tissue sources bear the same effect on the lymphatic junctions.

### Fibroblasts show tissue-specific effects on lymphatic junctions and permeability

To investigate tissue-specific effects of fibroblasts on lymphatic functions, we exposed hLECs to secretome from normal human lung fibroblasts (NHLF). While we found a significant increase in junction integrity and TEER with NHDF secretomes, we detected the opposite trend with NHLFs. There was a significant decrease in TEER from 87 ± 5 Ω.cm^2^ in the hLECs alone control to 72 ± 2 Ω.cm^2^ following treatment with NHLF secretomes (**Figure 8A**). Immunofluorescence imaging revealed an 18% reduction in ZO-1 expression following treatment with NHLF secretome, compared to hLEC alone (**Figure 8B and 8C**). We found no significant change in VE-cadherin expression compared to the control. Consistent with the reduction in TEER and ZO-1 expression, we observed a 2-fold increase in FD4 transport across hLECs treated with NHLF secretome compared to hLECs alone (**Figure 8D**). Despite differences in transport and TEER, we saw no effect of NHLFs on cell junction morphology between hLECs (**Supplementary Figure 3**). The opposing effects of lung and dermal fibroblasts demonstrate tissue-specific programming of lymphatic barrier function, which may underlie the varied susceptibility of different vascular beds to edema and inflammation.

**Figure 8.**
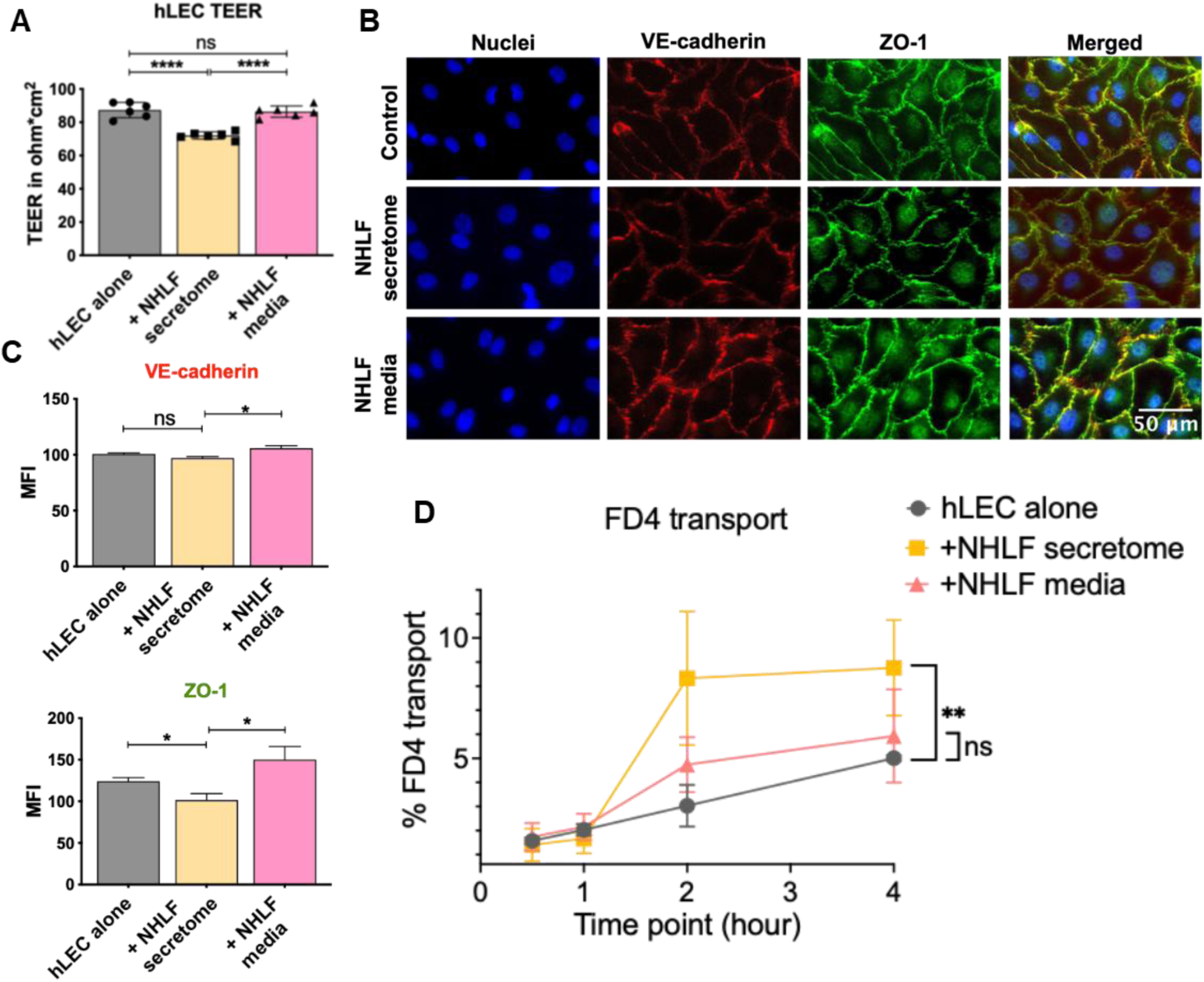
NHLF secretomes impair hLEC barrier function. **(A)** Transendothelial electrical resistance (TEER) measurements of hLEC monolayers following 48 hours of treatment with NHLF secretome or NHLF culture medium, compared with hLECs cultured alone. **(B)** Representative immunofluorescence images of VE-cadherin (red) and ZO-1 (green) in hLECs following 48 hours of treatment with NHLF secretome or NHLF culture medium. Nuclei were counterstained with Hoechst (blue). Scale bar = 50 μm. **(C)** Quantification of VE-cadherin and ZO-1 fluorescence intensity using ImageJ (FIJI). **(D)** FITC-dextran (4 kDa) transport across hLEC monolayers following treatment with NHLF secretome or NHLF culture medium. Statistical significance in panels A and C was assessed using one-way ANOVA followed by Dunnett’s multiple-comparisons test and Welch’s t-test (n=6). For the permeability assay (panel D), statistical significance was determined using two-way ANOVA followed by an appropriate post hoc multiple-comparisons test (n = 3). ns, not significant (*P* ≥ 0.05); *P* < 0.05 (\**), P < 0.01 (*\*\**), P < 0.001 (*\*\*\*), and *P* < 0.0001 (****). Images were adjusted to improve visualization. Quantitative analyses were performed using the original unprocessed images.

## DISCUSSION

Our findings identify stromal fibroblasts as active paracrine regulators of lymphatic endothelial barrier function through coordinated structural, functional, and transcriptional mechanisms (**Figure 9**). Integrating our experimental and transcriptomic data, the outcomes support a model in which stromal-endothelial crosstalk is a central determinant of lymphatic barrier homeostasis. Below, we discuss each of these mechanisms and their biological implications in detail.

**Figure 9.**
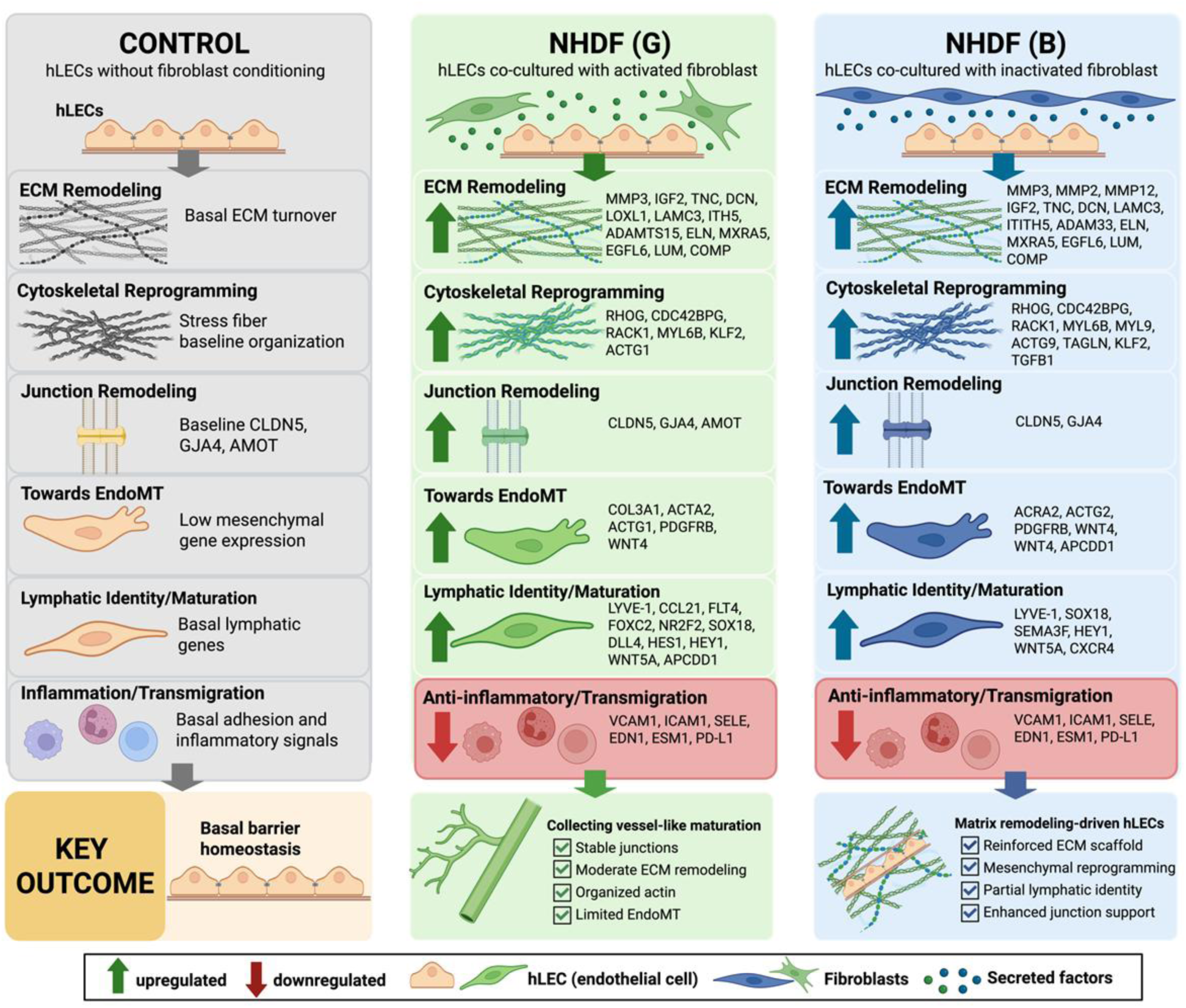
Schematic summary of the RNA-seq analysis of hLECs following 48 hours of co-culture with NHDF(G) or NHDF(B). Compared with hLECs cultured alone, NHDF(G) promoted lymphatic maturation, junctional remodeling, cytoskeletal organization, and moderate ECM remodeling, resulting in a collecting vessel-like phenotype. NHDF(B) preferentially induced ECM remodeling, mesenchymal reprogramming and cytoskeletal organization with partial lymphatic maturation. Fibroblasts were removed before transcriptomic analysis, and all RNA-seq data were generated from hLECs.

Dermal fibroblasts promote increased transendothelial electrical resistance (TEER), reduced 4 kDa dextran flux, and promote more continuous junctional localization of ZO-1 and VE-cadherin. All this indicates lymphatic barrier tightening through junctional reinforcement, consistent with the low-permeability, collecting-vessel phenotype required for efficient lymph propulsion. These data place fibroblasts alongside inflammatory and mechanical cues as determinants of lymphatic barrier properties and suggest that stromal-endothelial crosstalk is a key contributor to tissue-specific lymphatic homeostasis, edema resistance, and transport function.

The main objective of this study was to define how fibroblasts regulate lymphatic permeability and barrier function under homeostatic conditions. We demonstrate that dermal fibroblast secretomes are sufficient to decrease lymphatic endothelial permeability and strengthen junctional organization. Our data demonstrate that secretome treatment increases the expression of both tight and adherens junctions (ZO-1, Claudin-5, and VE-cadherin) on hLECs, compared to untreated controls. This interpretation is consistent with prior work showing that VE-cadherin is a central organizer of endothelial barrier integrity and that collecting lymphatic permeability is strongly regulated by VE-cadherin-dependent junctional architecture [2, 19, 20]. Additionally, researchers showed that 3 kDa dextran transport increases when VE-cadherin is blocked, suggesting its critical role in maintaining lymphatic barrier functions [1]. Our data also indicated a similar effect, where increased expression of VE-cadherin and ZO-1 at the junctions was associated with reduced dextran transport across the monolayer. The reduction in 4 kDa dextran transport that we observed upon treatment with fibroblast secretomes therefore provides a functional correlation of the junctional phenotype and supports the idea that fibroblast-derived soluble cues bias hLECs toward a low-permeability state.

Our co-culture data corroborated the findings from secretome-treated hLECs by demonstrating that the fibroblasts-induced barrier phenotype is maintained in a dynamic multicellular context and is accompanied by a shift in junction presentation of ZO-1 and VE-cadherin toward a continuous topology. Researchers showed that Human Umbilical Vein Endothelial Cells (HUVECs) co-cultured with neonatal foreskin fibroblasts had 50% reduced permeability to 70 kDa dextran [21]. Another study co-culturing HUVECs and lung fibroblasts showed a drop in dextran permeability for HUVECs from 61% to 39% after 7 days and to 7% after two weeks [22]. Consistent with these findings, our results show that dermal fibroblasts significantly reduce lymphatic permeability in a non-contact coculture system, suggesting that paracrine signals from fibroblasts promote lymphatic collecting vessel phenotypes. The pronounced increase in TEER, reduction in 4 kDa dextran flux, and shift toward more continuous ZO-1-positive junctions support the interpretation that dermal fibroblasts bias hLECs toward a collecting-vessel-like low-permeability phenotype, although additional in vivo work will be required to determine whether fibroblasts drive a full capillary-to-collecting phenotypic transition.

A major observation was the marked redistribution of ZO-1 junctions from discontinuous punctate to continuous junctions. Because continuous junctional organization is a hallmark of stabilized endothelial barriers, this result provides structural evidence that fibroblasts influence barrier function at the level of junction assembly and maintenance rather than simply altering marker expression. The fact that both NHDF(G) and NHDF(B) enhanced continuous ZO-1 presentation suggests that fibroblasts share a core barrier-supportive program even though their broader transcriptional effects diverge. Barrier strength is not determined solely by total junctional protein abundance. For example, researchers reported rat LECs had increased permeability and TEER after exposure to ethanol, despite no changes in ZO-1 or VE-cadherin expression or localization [23]. We observed that the overall expression of junction proteins, quantified using immunoblotting and RT-qPCR analyses, revealed no significant difference in the overall protein and mRNA expression of ZO-1 and VE-cadherin despite clear changes in junctional continuity and barrier function. This dissociation supports a model in which fibroblasts strengthen lymphatic barrier function by promoting junctional redistribution and stabilization of pre-existing proteins, potentially through altered cytoskeletal coupling, VE-cadherin complex organization, and cortical actin support, rather than by simply increasing total protein abundance. Our data further demonstrate that fibroblasts regulate lymphatic barrier physiology in parallel with substantial remodeling of the actin cytoskeleton. In particular, NHDF(G) conditioning induced a pronounced reduction in actin anisotropy, accompanied by the loss of aligned central stress fibers and redistribution of F-actin toward cortical and junction-associated regions. This pattern is consistent with established endothelial barrier models in which contractile stress fibers and centripetal tension promote junction destabilization and barrier disruption, whereas cortical actin supports cadherin-mediated junction stabilization and enhanced barrier integrity [24]. The more pronounced effect of NHDF(G) compared to NHDF(B) further suggests that fibroblast activation state modulates the magnitude of endothelial cytoskeletal remodeling. Collectively, these findings provide a physiologically plausible structural basis for the observed increase in TEER and decrease in dextran permeability. They suggest that fibroblasts enhance lymphatic barrier function, at least in part, by promoting a shift from a tension-generating stress fiber network toward a junction-supportive cortical actin architecture.

To define the molecular mechanisms underlying lymphatic responses to fibroblast stimulation, we performed bulk RNA sequencing and analyzed differentially gene expression. Recent transcriptomic studies have uncovered the remarkable heterogeneity of LECs, revealing that these cells serve functions beyond fluid transport and immune cell trafficking - they also participate in diverse biological processes and exhibit dynamic interactions with other cell types [25, 26]. A single-cell RNA-seq study identified six distinct LEC subtypes in human lymph nodes [27]. Here, we identify two distinct fibroblast-driven programs that promote endothelial remodeling and junctional integrity but differ fundamentally in their molecular imprints. Co-culture with NHDF(G) induced a transcriptional landscape consistent with reinforcement of lymphatic identity and junctional integrity. The upregulation of canonical lymphatic regulators such as SOX18, NR2F2 and mature lymphatic markers including CCL21 and VEGFR3 suggests that NHDF(G) provide cues that sustain or enhance lineage specification. Importantly, this was accompanied by increased expression of junction-associated genes (CLDN5, GJA4, AMOT) and cytoskeletal regulators (Rho GTPases, ACTG1, CDC42BPG), indicating coordinated remodeling of both structural and mechanical components of the endothelial barrier. AMOT is recognized as a key linker between actin filaments and junctional complexes, and its upregulation supports a model in which cytoskeletal reorganization is closely coupled to junction stabilization. The concurrent enrichment of mechanosensitive and signaling mediators such as KLF2, TGFB1, and ERK/Notch pathways further suggests that NHDF(G) may promote a flow-responsive, dynamically adaptive endothelial phenotype. The simultaneous downregulation of VCAM1, ICAM1, and SELE further supports a more quiescent and less inflammatory endothelial state, which aligns well with the increased TEER, reduced dextran flux, and more continuous junction morphology observed experimentally. This is consistent with the physiological role of lymphatic vessels in maintaining immune homeostasis while limiting excessive leukocyte infiltration (collecting-vessel type).

In contrast, NHDF(B) co-culture elicited a noticeably different transcriptional hierarchy dominated by ECM remodeling and mesenchymal-like activation. While some overlap with NHDF(G)-induced pathways was evident, the scale and preferential upregulation of ECM-associated genes, including MMPs, POSTN, LOX, and laminins, suggest a microenvironment defined by dynamic matrix remodeling and altered biomechanical properties. The coordinated induction of ACTA2, PDGFRB, and WNT-related genes suggests that NHDF(B) biases hLECs toward partial EndoMT-like remodeling rather than toward a purely barrier-stabilizing phenotype. Because WNT, p38 MAPK, and ECM-remodeling programs are commonly associated with fibrosis, endothelial plasticity, and altered biomechanical signaling, this profile suggests that NHDF(B) may establish a more remodeling-permissive microenvironment. Notably, NHDF(B) still preserved some junctional-supportive features, indicating that barrier reinforcement and remodeling are not mutually exclusive states, but the balance of pathways was clearly shifted away from the more quiescent, collecting-vessel-like program induced by NHDF(G).

Direct comparison between NHDF(G) and NHDF(B) conditions highlights two mechanistically distinct modes of fibroblast-hLEC interaction. While a shared core of regulated genes indicates common responses to fibroblast-derived signals, divergence in condition-specific gene sets reflects specialization toward either vascular stabilization (NHDF(G)) or matrix remodeling and transition (NHDF(B)). Enrichment of lymphatic maturation and immune-regulatory genes in NHDF(G) suggests a supportive niche that reinforces endothelial function, whereas the prominence of ECM- and WNT-related pathways in NHDF(B) is consistent with a pro-remodeling, potentially pathological microenvironment. These findings support a model in which fibroblast heterogeneity dictates endothelial fate by modulating the balance between cytoskeletal organization, junctional integrity, and matrix interactions. Thus, our RNA-seq study reveals that dermal fibroblasts are not passive structural components but active regulators of endothelial phenotype, capable of reprogramming endothelial junctional architecture through distinct molecular routes.

This in vitro system provides a tractable platform to dissect stromal-endothelial signaling and establishes key principles of fibroblast-mediated lymphatic barrier regulation. Building on this foundation, future work should systematically map donor-to-donor variation by expanding both LEC and fibroblast sources to capture the breadth of human stromal diversity. Next-generation models that incorporate luminal and transmural flow, cyclic mechanical strain, and 3D vessel architecture will be important for defining how fibroblast-derived cues integrate with mechanical forces to tune lymphatic junction dynamics and permeability. Introducing additional perivascular and immune cell populations, including smooth muscle cells and leukocytes, will be critical to understand how multicellular microenvironments shape lymphatic barrier tone in homeostasis and inflammation. In a complementary experiment, lung fibroblast secretomes promote a more permeable and destabilized hLEC junctional profile, in contrast to the dermal fibroblast. Extending these studies to fibroblasts from other organs such as cardiac, mesenteric, and renal tissues, and pairing tissue-matched fibroblast-LEC combinations will help define organ-specific stromal programs. Finally, targeted perturbation and profiling of fibroblast secretomes will be essential to identify the specific mediators and pathways responsible for the distinct barrier phenotypes described here.

Taken together, this work shows that fibroblasts are key regulators of lymphatic endothelial barrier function in the local microenvironment, where stromal and immune cells shape LEC behavior. Dermal fibroblasts enhance LEC integrity by strengthening tight and adherens junctions and remodeling the ECM, thereby limiting paracellular transport, while fibroblast heterogeneity and tissue origin generate distinct barrier phenotypes that warrant further investigation. These findings have important implications for diseases with lymphatic dysfunction, including cardiovascular conditions, where altered fibroblast states may weaken lymphatic integrity, impair drainage, and worsen tissue congestion. In this context, our thrombin data illustrate how inflammatory mediators elevated in pathology can further compromise fibroblast-conditioned barriers, underscoring the need to define stromal and inflammatory pathways that can be targeted to preserve or restore lymphatic function.

## ACKNOWLEDGMENTS

Financial support was provided by the NIGMS MIRA 1R35GM142835-01(KM), as well as the GEM fellowship (AA) and Drs. Wayne T. & Mary T. Hockmeyer Summer Fellowship Award from the Biological Sciences Ph.D. Program (AA). We would also like to acknowledge Michele Kaluzienski for copy-editing the finalized document, the BioWorkshop Core facility, and the Stroka Lab for helping us establish the JAnaP analysis in our lab.

## MATERIALS AND METHODS

### Cell culture and co-culture model

Human primary dermal lymphatic endothelial cells (hLECs) (PromoCell, C-12217), isolated from adult skin (experiments are representative of data from n=8 donor cells) were seeded onto the bottom side of Transwell inserts (1 μm pore size, Falcon®, 353103), coated with 50 μg/mL rat tail collagen I (Corning, 354236). After seeding hLECs for 1 hour at 37 °C, inserts were turned over and cells were allowed to grow for 5 days in Endothelial Cell Growth Medium MV2 (EGM-V2; PromoCell, C-22121) until the hLECs formed a confluent monolayer. Monolayer formation was confirmed by measurement of transendothelial resistance (TEER) or by staining with CellMask™ Plasma Membrane Stain (ThermoFisher, C10046).

Normal human primary dermal fibroblasts (NHDF) derived from adult human skin, which represent a mixed population of papillary and reticular fibroblasts (experiments are representative of data from n=3 donor cells) were cultured in two conditions: (1) NHDF(G) (PromoCell, C-12302) in Fibroblast Growth Medium-2 (PromoCell, C-23120) containing 2% FBS, 1 ng/mL fibroblast growth factor, and 5 μg/mL insulin, or (2) NHDF(B) in fibroblast basal medium without supplements. For secretome collection, fibroblasts were cultured with EGMV2 for 24 hours. Media was then collected and centrifuged to remove debris, and the resulting conditioned medium (secretome) was used for treatment. Normal human primary lung fibroblasts (NHLF; Lonza, CC-2512) isolated from adult lung tissue (experiments are representative of data from n=2 donor cells) were cultured in FGM-2 (Lonza, CC-3132) and processed similarly to NHDFs for secretome collection. The hLEC monolayers on inserts were conditioned at the top side with either (1) NHDF cells [NHDF(G) or NHDF(B)] or (2) their corresponding secretomes, for 48 hours. Following co-culture, secretomes were removed, and fibroblasts were detached from inserts using a sterile cotton applicator (Dealmed, B074MJHSGG). hLECs were then processed for other assays. For thrombin experiments, 2 U/mL thrombin was added to the hLEC monolayer at the top. After treatment, inserts were washed thoroughly, and hLECs were co-cultured with NHDF(G) cells for 48 hours prior to downstream analysis.

### Transendothelial electrical resistance (TEER)

To assess the confluence and barrier integrity of hLEC monolayers, TEER was measured using a Millicell® ERS-2 Voltohmmeter (Millipore). Electrodes were carefully positioned in both the upper and lower chambers of the Transwell system, ensuring contact with the culture medium. Initial resistance values (in ohms) were recorded and corrected by subtracting background resistance values obtained from blank inserts (without cells). Final TEER values (Ω·cm²) were calculated by multiplying the resistance of the cell-containing inserts by the surface area of the 12-well Transwell membrane (1.12 cm²). These measurements were used to confirm monolayer confluency and hLEC barrier integrity prior to downstream assays.

### In vitro lymphatic transport assay

To evaluate the effect of fibroblasts or their secretomes on endothelial permeability, hLEC monolayers were subjected to a transport assay following co-culture or treatment [28]. After removing fibroblasts or secretomes, hLECs were washed and incubated with fresh EGMV2 basal media. Media containing 20 µg/mL FITC-labeled dextran 4 kDa (FD4; Millipore Sigma, 46944) was added to the upper chamber of the Transwell insert. The transport assay was carried out for 8 hours at 37 °C. At defined timepoints, aliquots were collected from the lower chamber (plate bottom), and fluorescence intensity was measured using a microplate reader (Tecan Spark) and standard curve. The amount of FD4 transported across the hLEC monolayer was used as a quantitative readout of lymphatic permeability.

### Immunofluorescence staining, confocal microscopy and imaging analyses

For immunofluorescence imaging and downstream analysis, hLECs were fixed in 2% paraformaldehyde (PFA) in 1X phosphate-buffered saline (PBS; Thermo Fisher, J19943.K2) for 15 minutes at room temperature, then permeabilized with 0.1% Triton X-100. Cells were blocked in 2% fetal bovine serum (FBS; Gibco, 10100-147) in PBS for 1 hour. Membranes were carefully excised and mounted onto glass slides. Cells were incubated overnight at 4°C with primary antibodies: mouse anti-VE-cadherin (1:200; BD Biosciences, 55566) and rabbit anti-ZO-1 (1:200; Invitrogen, 33-9100). The following day, cells were incubated with species-specific Alexa Fluor-conjugated secondary antibodies-donkey anti-mouse Alexa Fluor 488, 555, or 647 (1:400; Invitrogen, A21202, A31570 or A31571) and donkey anti-rabbit Alexa Fluor 488, 555, or 647 (1:400; Invitrogen, A32790, 31572 and A31573) for 2 hours at room temperature. Washes were performed with 0.1% Tween-20 in 1X PBS (BioShop, TWN510) before and after antibody incubations. Nuclei were counterstained using Hoechst 33342 (1:10,000) and cover slips were mounted using VECTAshield Antifade Mounting Medium (Vector Laboratories, H-1200-10). Images were acquired using a Zeiss Observer 7 fluorescence microscope (20X/63X objectives) or an FV3000 laser scanning confocal microscope (20X/100X objectives).

### Immunoblot assay and qPCR

After co-culture, hLECs were lysed using RIPA lysis buffer (Thermo Fisher, 89900). The resulting cell lysates were subjected to gel electrophoresis and immunoblotting as previously described [29]. Briefly, 20 μg of protein per lane was denatured in Laemmli buffer and separated on a 4–20% SDS-PAGE gel (Mini-Protein TGX, BioRad) at 110V for 1 hour. Proteins were then transferred to nitrocellulose membranes (Trans-Blot Turbo, BioRad) using a semi-dry transfer system (1.5 mA, 25 V for 7 minutes), and membrane transfer was confirmed via Ponceau S staining. The membranes were sectioned into strips, blocked in 2% BSA (prepared in TBS-T) for 1 hour, and incubated overnight at 4°C with primary antibodies: mouse anti-VE-cadherin (BD Biosciences, 55566) and rabbit anti-ZO-1 (Invitrogen, 33-9100) at a 1:500 dilution. After washing, the membranes were incubated for 1 hour at room temperature with secondary antibodies: IRDye® 680RD donkey anti-Rabbit, donkey anti-Mouse, and IRDye® 800CW donkey anti-Rabbit (all at 1:20,000 dilution). Protein bands were visualized using the Odyssey® Imaging System (LI-COR).

To quantify expression of genes encoding adherens and tight junction proteins, total RNA was isolated from hLEC samples using TRIzol™ reagent (Invitrogen, 15596026), following the manufacturer’s protocol. To obtain sufficient RNA for analysis four transwell inserts were used as technical replicates, and RNA from these inserts were pooled for each biological replicate. A total of 500 ng of RNA per sample was reverse transcribed into cDNA using the High-Capacity cDNA Reverse Transcription Kit (Applied Biosystems, 4374966), according to the manufacturer’s instructions. Quantitative real-time PCR (RT-qPCR) was performed using TaqMan™ Gene Expression Assays specific for VE-cadherin (CDH5, Hs00901465), ZO-1 (TJP1, Hs01551871), PROX1 (Hs00896293), LYVE-1 (Hs00272659), and the housekeeping gene GAPDH (Hs02786624). Relative gene expression levels of hLECs after co-culture were calculated using the comparative Ct method (ΔΔCt) and relative fold changes (2^^−ΔΔCt^) were determined with respect to control.

### Image Processing

Image analysis was conducted using FIJI (ImageJ2, version 2.14). For cell count, we calculated the number of cells in one frame and multiplied this with total transwell area. For quantification of mean fluorescence intensity (MFI) of proteins, threshold were set to calculate only the junctional staining. For detailed single-cell analysis, the Junction Analyzer Program (JANaP) was utilized, as previously described [17]. Junction morphology of hLECs was classified as either continuous (linear or mature) or discontinuous, which included isolated, jagged, punctate, or perpendicular junctional patterns. In addition, cell shape descriptors such as circularity and solidity were quantified to assess morphological changes.

F-actin alignment was assessed using the FibrilTool plugin in ImageJ (FIJI) as described by Boudaoud et al. [30]. Quantification was performed on immunofluorescence images within regions of interest (ROIs) for central stress fibers, and anisotropy values were extracted. Anisotropy values ranging from 0 (random organization) to 1 (perfect alignment) indicate increasing degrees of actin fiber alignment. At least 15 cells per condition were analyzed across three independent biological replicates.

### Differential RNA gene expression analysis

Total RNA was extracted from three groups of hLECs, (1) without co-culture (hLECs alone), (2) co-cultured with NHDF(G), and (3) co-cultured with NHDF(B) using TRIzol™ reagent (Invitrogen, 15596026). RNA quality was initially assessed using the NanoDrop™ 2000c spectrophotometer (Thermo Scientific). Approximately 2 µg of RNA from four biological replicates per group were sent to Genewiz, Azenta Life Sciences, for sequencing. Briefly, RNA integrity was evaluated, ensuring RIN values exceeded 9.0. RNA sequencing was performed on an Illumina platform with paired-end reads (2×150 bp), generating approximately 50 million reads per sample.

In the bioinformatics pipeline, raw reads were trimmed using Trimmomatic v0.36 and then aligned to the GRCh38 human reference genome using STAR aligner v2.5.2b. Differentially expressed genes (DEGs) were identified using PyDESeq2 algorithm (pydeseq2 v0.5.2), with p-values and log₂ fold changes calculated via the Wald test. Genes with an adjusted p-value < 0.05 and an absolute log₂ fold change > 1 were considered significantly differentially expressed. Volcano plots depicted the overall distribution of up- and downregulated genes. Gene set enrichment analysis (GSEA, gseapy v1.1.10) was then performed using the gene sets using the molecular signature (MSigDB) database: Gene Ontology Biological Process (GOBP v2025.1) gene sets.

### Statistical Analysis

All analysis was performed using GraphPad software (version 10.4.2), and data are represented as mean ± standard error of the mean (SEM). Statistical analysis was performed using one-way ANOVA and unpaired two-tailed *t*-test; *p* > 0.05 = not significant (ns), *p* ≤ 0.0001 = extremely significant (****). For the permeability (transport) assay, statistical significance was determined by two-way ANOVA followed by appropriate post hoc test for multiple comparisons. ns, not significant (*P* ≥ 0.05); *P* < 0.05 (\**), P < 0.01 (*\*\**), P < 0.001 (*\*\*\*), and *P* < 0.0001 (****).

