## Supplement for "Fibroblasts dynamically regulate lymphatic barrier function by modulating cell-cell junctions"

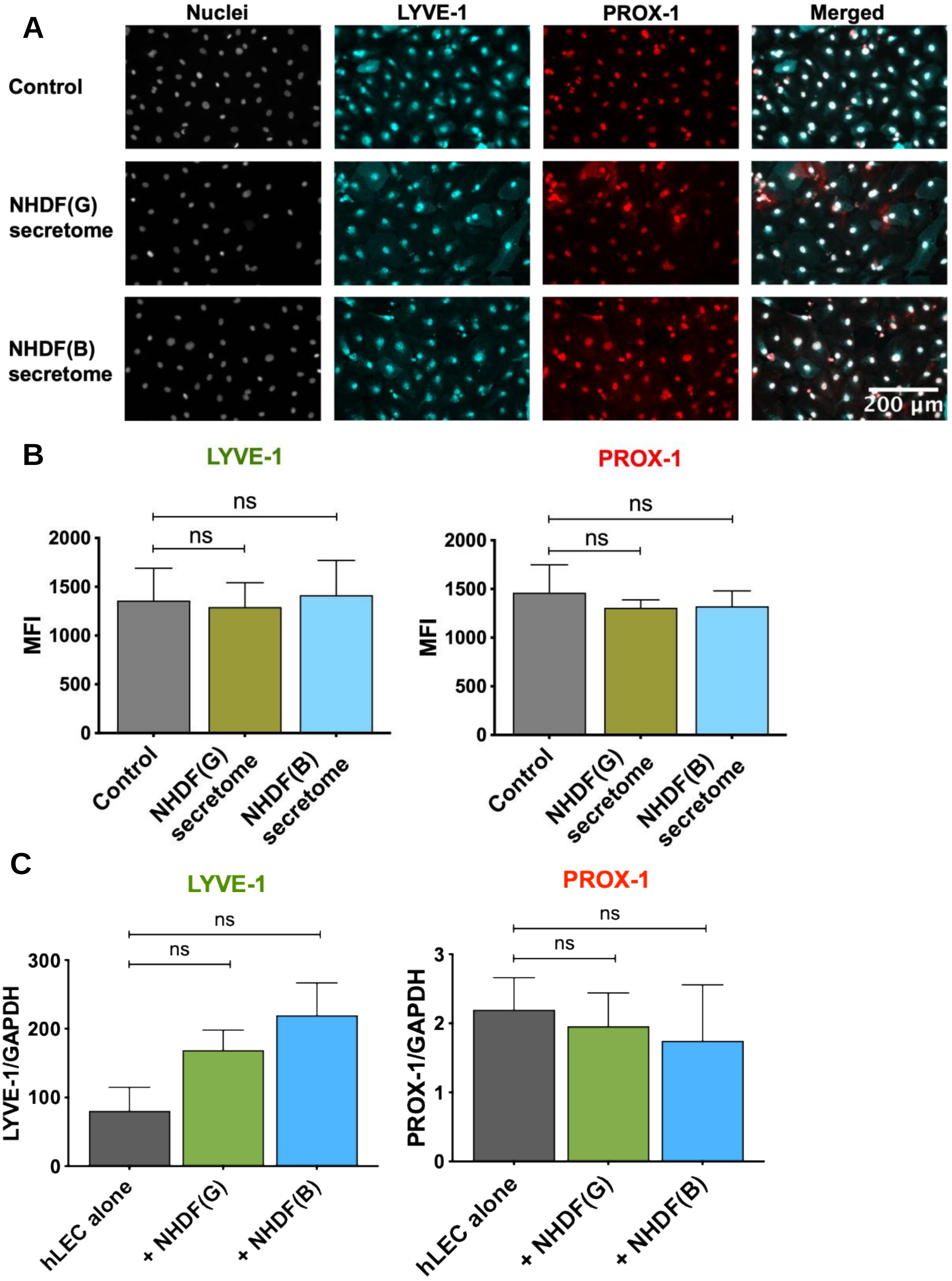


**Supplementary Figure 1. (A)** Representative immunofluorescence images of PROX1 and LYVE-1 on hLECs following treatment with NHDF secretomes. **(B)** Quantification LYVE-1 and PROX1 expression in hLEC monolayers treated with NHDF secretomes. (**C**) RT-qPCR analysis of mRNA expression levels of LYVE-1 and PROX1 in hLECs following NHDFs co-culture.

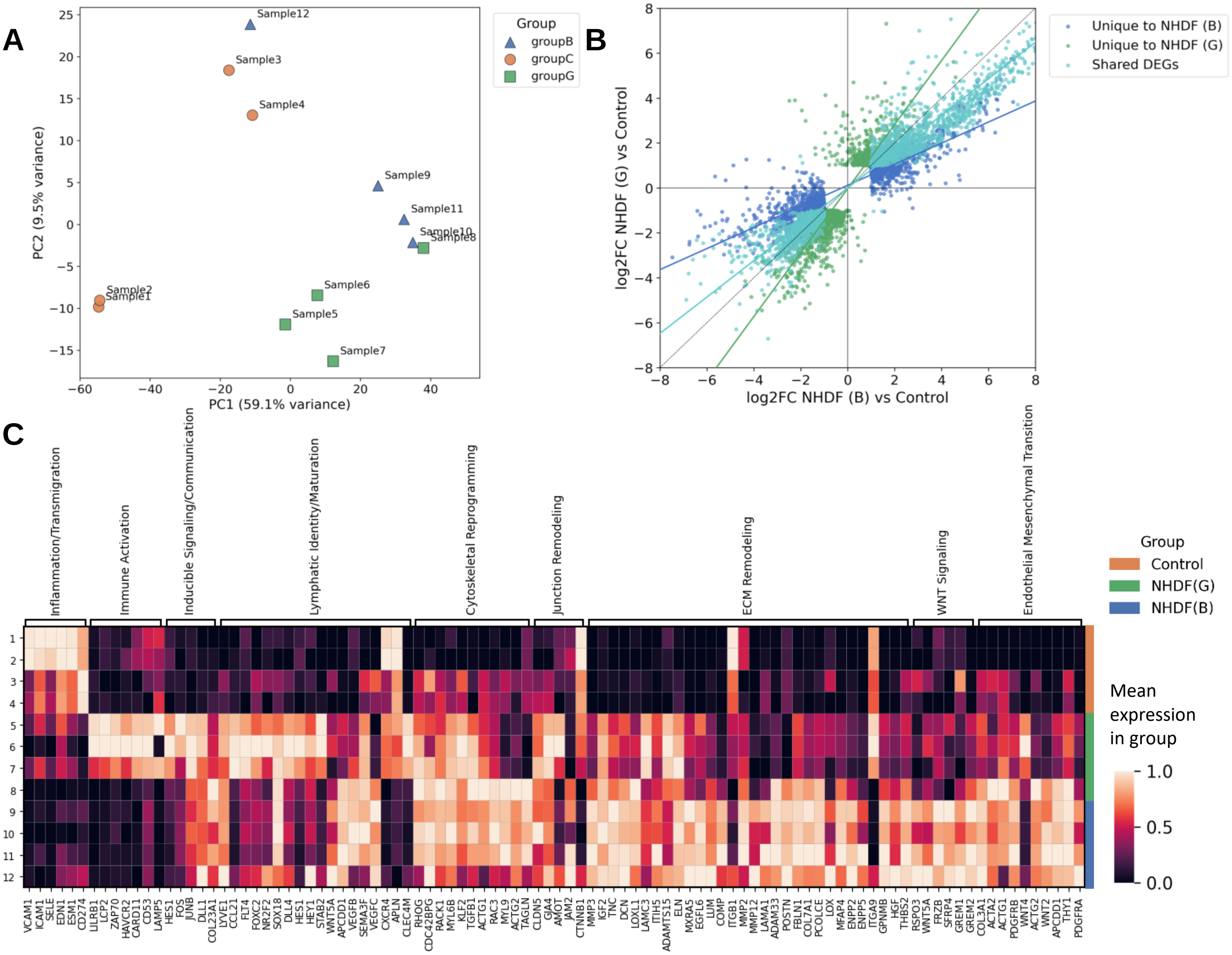


**Supplementary Figure 2. Expanded sample PCA and representative genes associated with fibroblast-induced transcriptomic remodeling of hLECs. (A)** Per-sample comparison of Principal Components 1 and 2 indicating sample similarity. **(B)** Comparison of $\log_2$ fold change of NHDF (G) vs Control (y-axis) and NHDF (B) vs Control (x-axis) within significantly differentially-expressed genes (DEGs). **(C)** Normalized expression of selected genes grouped by major biological processes in hLECs cultured alone (Control) or following 48 hours of co-culture with NHDF(G) or NHDF(B). Genes were selected to represent broad biological processes. Color intensity represents relative expression levels across samples.

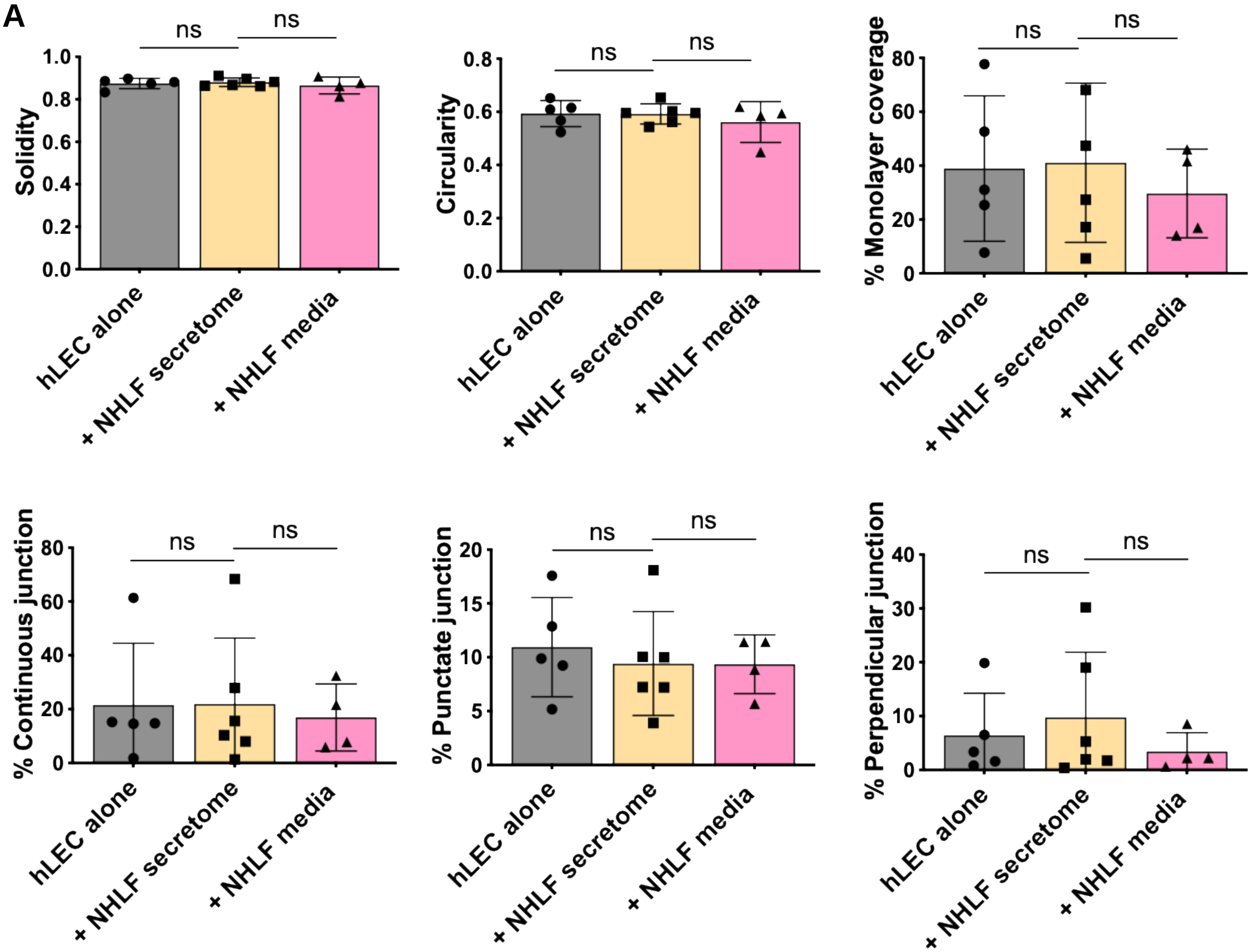


**Supplementary Figure 3.** Quantitative analysis of ZO-1 distribution and organization in hLEC monolayers using junction analyzer program, JAnaP after 48 hours of co-culture with NHLF secretomes.
